# Longitudinal EEG identification using sensor-level phase and complex coherency across sessions and functional states

**DOI:** 10.64898/2026.10.26.761764

**Authors:** Dmitry Goldstein, Anthonios Deeb, Bashar Nassir, Saleh Sawaed, Nezer Jacob Zaidenberg

**Author notes:** Corresponding author: Dmitry Goldstein.

## Abstract

Sensor-level EEG coherency can distinguish individuals, although its phase and magnitude patterns may reflect neural activity, source mixing, and recording geometry. We evaluated two geometric representations of complex coherency—unit phasors and full complex coherency— alongside triangle holonomies and magnitude-only descriptors. Using direct enrollment–probe matching without a trained identity classifier, we analyzed three independent longitudinal cohorts: RestCog (*N* = 60; approximately one month), L-FAME (*N* = 44; six weeks), and Dortmund (*N* = 206; approximatelyfive years). In eyes-closed rest-to-rest matching, alpha phasors identified 81.67% of participants in RestCog (AUC 0.9473) and 79.55% in L-FAME (AUC 0.9735); full complex coherency identified 78.33% and 84.09%, respectively. After five years in Dortmund,fixed row-standardized alpha–beta fusion yielded 77.67% rank-1 identification (AUC 0.9744) for phasors and 70.39% (AUC 0.9527) for full coherency. Changes in functional state exposed different strengths: full coherency preserved higher rank-1 accuracy in some monthly RestCog rest-to-task comparisons, whereas alpha phasors retained higher verification AUC. Switching between eyes-closed and eyes-open recordings in Dortmund substantially weakened identification with both representations. Triangle holonomies also retained individual information across visits. An exact factorization of complex cosine shows how magnitude-pattern alignment and magnitude-weighted phase agreement contribute to full-coherency matching. Together, the results establish longitudinally discriminative structure in measured scalp EEG coherency over weeks to years, while distinguishing persistence across visits from robustness to changes in recording state.

## 1 Introduction

Longitudinal EEG identification requires a pattern that remains recognizable when the same person returns for another recording, potentially under different conditions. Functional MRI connectivity profiles can distinguish individuals across scans [1, 5], and resting-state EEG connectivity can likewise identify participants [21]. Yet identification alone does not reveal which aspects of electrophysiological connectivity remain distinctive across visits, or how their representation affects recognition.

Time and functional state pose different challenges. An EEG pattern may be reproducible during repeated rest but change during mental arithmetic, memory recall, or eye opening. Conversely, a feature that distinguishes individuals across tasks within a single visit may not persist for months or years. Previous studies have examined task-related variability in EEG connectivityfingerprints [6], their temporal robustness [3], and longitudinal individual differences in resting EEG dynamics [9]. The reliability of connectivity and network measures also varies with the descriptor and recording condition [4]. Recent EEG-identification studies have also exploited network structure and learned representations: Li et al. [13] incorporated the spatial arrangement of scalp channels into functional-connectivity graphs, Tian et al. [20] combined multiband phase-locking-value graphs with a graph convolutional network, and Oztemel and Soysal [15] evaluated attention-based autoencoder features across visits separated by ten days. These approaches motivate the study of person-specific sensor-level EEG structure, but use different representations, training procedures, or recording protocols. Here we examine persistence across visits and transfer across states as related, but separate, properties, focusing on the signed phases and full complex entries of coherency without learning an identity classifier.

Recent studies using the same public cohorts highlight the distinction between these questions. A preprint based on RestCog assessed both test–retest reliability and subject identifiability of frozen EEG representations [7]. In Dortmund, spectral biomarker candidates have been evaluated for five-year stability and eyes-open/eyes-closed sensitivity [16]. The original L-FAME benchmark instead addresses meditation-state and technique decoding, along with cross-session model adaptation [12]. These investigations provide relevant context, but their reported outcomes are not matched enrollment–probe identification baselines.

An important interpretive boundary arises from recording at the scalp. Each electrode samples a mixture of signals influenced by volume conduction, head anatomy, sensor placement, and the EEG reference. Signed cross-spectral phase may change under such mixing; a pattern that identifies participants therefore need not represent stable coupling between cortical sources. Throughout this study, an EEG fingerprint means an individually discriminative pattern in the recorded sensor-space signals, without a claim about its uniquely neural origin [2, 14].

Complex coherency offers a way to ask which features carry this individual information. For channels *i* and *j*, the frequency-domain relationship can be written as 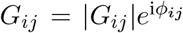: the magnitude measures a normalized cross-spectral association at the sensors, while the signed angle *ϕ*_*ij*_ describes their relative phase. Phase synchronization has a long history in EEG analysis [11]; imaginary coherency and phase-lag measures were developed in part to address misleading coupling associated with instantaneous mixing and common sources [14, 18]. Here we retain the signed phase of each coherency edge and compare the resulting pattern across recordings, rather than reducing coherency to its magnitude or to a real-valued phase-lag summary.

The complex representation also leads naturally to two distinct matching geometries. Mapping each edge phase to a unit phasor, 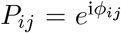, places the *m* = *p*(*p* − 1)*/*2 independent edge phases of a *p*-channel recording on a product of circles, (*S*^1^)^*m*^. At the final matching stage, phasor similarity gives equal weight to each edge’s phase agreement. Matching the full complex entries of *G*, by contrast, retains the magnitudes: strong edges in both recordings contribute more to phase agreement, and the alignment of magnitude patterns affects the normalized similarity. The two scores therefore emphasize different aspects of the same coherency matrix and may rank candidate identities differently.

We also examine what remains of the phase fingerprint after removing node-wise phase freedom. Under the transformation *ϕ*_*ij*_ *↦ϕ*_*ij*_ +*θ*_*i*_ −*θ*_*j*_, the phase accumulated around a triangle is unchanged. These triangle holonomies provide a gauge-invariant representation of the edge-phase pattern, related to the geometry of angular synchronization [17]. Comparing holonomies across visits tests whether individual information persists after this algebraic reduction. Magnitude-only matching supplies a further comparison for interpreting the full complex score.

We apply these representations to three independent longitudinal EEG cohorts. In RestCog, the main comparison spans approximately one month, with a 90-minute within-cohort reference; the recordings include eyes-closed rest and cognitive conditions. Dortmund includes eyes-closed and eyes-open recordings collected approximately five years apart, while L-FAME compares rest and silent meditation across six weeks. Because the recording protocols differ, these intervals are not points along a single decay curve. Same-state enrollment–probe matching tests persistence across visits, whereas cross-state matching examines sensitivity to recording condition. Using direct matching without a trained identity classifier, we evaluate rank-based identification and verification AUC, alongsidefixed band-fusion and recording-duration comparisons. Our aim is to determine how phase-only and full complex matching preserve individual discriminability in sensor-level EEG, when their rankings differ, and what holonomy and magnitude reveal about these measured representations.

## 2 Materials and methods

### 2.1 EEG datasets and longitudinal study design

We evaluated individual EEG fingerprints in three independent, publicly available longitudinal cohorts. Each comparison enrolls participants at an earlier visit and matches their recordings to probes obtained at a later visit. Same-state comparisons examine persistence across visits; cross-state comparisons additionally test whether this persistence survives a change in functional condition. The cohorts have different recording protocols and are analyzed separately, without cross-dataset identity matching or a pooled estimate of temporal decay.

- **RestCog** (OpenNeuro ds004148; 22). This dataset contains three EEG sessions from 60 young adults (18–28 years), of which 59 common channels were retained. Session 01 is the enrollment visit; session 03 provides the principal probes approximately one month later, while session 02 supplies a secondary approximately 90-minute reference. Recordings include eyes-closed rest (EC), mental arithmetic (MA; serial subtraction), episodic memory (ME), and internally generated music (MU; silent singing). The three cognitive conditions are also eyes closed.
- **Dortmund Vital Study** (OpenNeuro ds005385; 8). The longitudinal analysis comprises 206 participants and 64 EEG channels, with visits separated by approximately five years. Pre-task eyes-closed (EC) and eyes-open (EO) recordings permit both same-state and cross-state comparisons. The source dataset reports 208 follow-up participants; 206 are included in the present paired analysis.
- **L-FAME** (dataset repository; 10, 12). Recordings were obtained before (`ses-premedita`) and after (`ses-posmedita`) a six-week meditation course. We analyzed the 44 participants with paired visits (the public dataset contains 74 participants at baseline) using a fixed panel of 55 EEG channels. The two retained eyes-closed conditions are the first resting block (CE01, `restCE01`) and silent meditation (`slMedita`); both same-state and cross-state comparisons are included. This study tests fingerprint persistence, not the causal effect of training.

The principal RestCog analysis enrolls participants in session-01 EC and probes them in session-03 EC, MA, ME, or MU. Thus its monthly cross-state series does *not* include MA-to-MA, ME-to-ME, MU-to-MU, or task-to-task enrollment. The secondary session-01-to-02 analysis comprises three EC/task 2 × 2 panels (EC/MA, EC/ME, and EC/MU); their shared EC-to-EC control is counted once, giving ten unique directed comparisons. Dortmund and L-FAME each contribute a complete 2 × 2 state panel. In every panel, the arrow denotes enrollment state at the earlier visit followed by probe state at the later visit; changing the state direction does not reverse chronological visit order.

**Table 1.** Longitudinal EEG cohorts and retained recording supports.

| Cohort / interval | *N* / channels | State comparisons | Support per recording |
|---|---|---|---|
| RestCog / 1 month | 60 / 59 | EC→EC, MA, ME, MU | 6 × 50 s = 300 s |
| RestCog / 90 min | 60 / 59 | EC/task 2 × 2 panels | 6 × 50 s = 300 s |
| Dortmund / 5 years | 206 / 64 | EC/EO 2 × 2 | 3 × 50 s = 150 s |
| L-FAME / 6 weeks | 44 / 55 | CE01/slMedita 2 × 2 | 1, 2, 3 × 50 s<br>= 50, 100, 150 s |

For the RestCog monthly EC duration analysis, an additional 3×50-s condition uses 150 s, while × 100 and 2 × 150 s are compared with 6 × 50 s at the same total 300-s support. Changing window length at afixed multitaper time–bandwidth product also changes spectral smoothing; this does not isolate the effect of window count. L-FAME uses thefirst one, two, or three contiguous 50-s windows (50, 100, or 150 s), and the complex coherency matrices are averaged before extracting phase or holonomy.

In L-FAME, nine raw channels (C4, P4, O1, O2, PO3, PO4, PO8, PO9, PO10) were excluded consistently before average referencing on the basis of pre-biometric raw-signal quality control, not identification performance. The retained research audit reported 44 paired participants and no fatal recording failures for the analyzed windows. The analysis hierarchy organizes completed comparisons; it is not claimed to have been prospectively preregistered. Session labels alone do not establish a controlled cap-removal or cap-reapplication intervention.

### 2.2 EEG preprocessing and spectral estimation

The archived RestCog/Dortmund pipeline restricts each cohort to a common, consistently ordered EEG montage, applies average reference, a 50-Hz notch, resampling to 128 Hz, a 0.5-Hz high-pass, and channel standardization. Analysis windows are non-overlapping. The exact standardization scope and temporal crop positions require confirmation from the archived run configurations. L-FAME follows the documented fixed representation and window-averaging scheme with the 55-channel panel; the run-specific preprocessing configuration still requires reconciliation. In particular, the L-FAME acquisition metadata specify 60-Hz mains frequency [10], whereas the inherited project pipeline records a 50-Hz notch. These dataset-level metadata do not establish the run-specific notch settings. Filter implementation and artifact-handling conventions require confirmation from the run configuration.

Within window *w*, spectral matrices are estimated using discrete prolate spheroidal sequence multitapers with time–half-bandwidth product *NW* = 3 and *K* = 5 tapers [19]. Writing 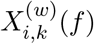 for the tapered Fourier coefficient of channel *i*,

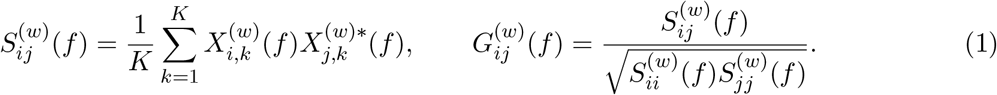

Coherency normalization precedes frequency averaging. For band *b*, the band-level matrix and recording template are

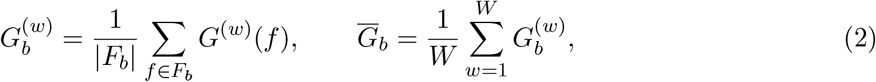

with alpha 8 ≤ *f <* 13 Hz and beta 13 ≤ *f <* 30 Hz. Frequency-wise normalization is not interchangeable with normalizing an already averaged cross-spectrum. Below, *G* denotes *G*_*b*_ unless otherwise specified. It is Hermitian; its diagonal is excluded from every primary matching score. Including identical unit diagonal entries would change the normalization and the scores.

The matrices are computed directly from scalp-electrode recordings, not reconstructed cortical sources. In an idealized instantaneous linear sensor mixture *y*(*t*) = *Mx*(*t*) with real mixing matrix *M*, the cross-spectrum transforms as *S*_*y*_(*f*) = *MS*_*x*_(*f*)*M* ^T^. Entrywise coherency normalization, and hence the resulting signed phases, are not generally invariant to this transformation. Accordingly, the matching descriptors are interpreted as sensor-level features; neither unit phasors nor triangle holonomies remove the effects of arbitrary volume conduction, sensor mixing, or re-referencing.

An independent Dortmund EC-to-EC baseline used alpha-filtered time-domain Pearson correlation matrices (206 participants, 64 channels, 3 ×50 s). The three fixed matrix-matching rules and preprocessing details are reported in the Supplementary Material. Because this archived control used a separate signal-processing pipeline, it is treated as a descriptive comparison rather than a preprocessing-matched ablation.

### 2.3 Phase and complex coherency geometry

The retained recording-level phase and phasor matrices are

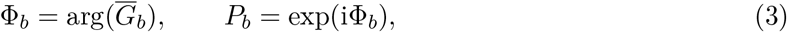

where exponential and argument are applied entrywise. Hermitian symmetry implies *ϕ*_*ji*_ = − *ϕ*_*ij*_ modulo 2*π*. The strict-upper phase vector belongs to (*S*^1^)^*m*^, with *m* = 1711 for RestCog, *m* = 2016 for Dortmund, and *m* = 1485 for L-FAME.

For recordings *A* and *B*, its extrinsic circular similarity is

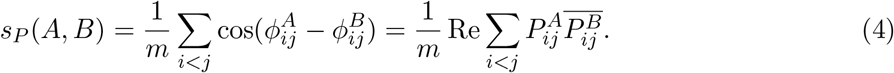

This score is periodic, lies in [−1, 1], and requires no phase unwrapping. It gives equal matching weight to every retained edge. It is equivalently a monotone transform of the squared chordal distance between unit-phasor vectors: 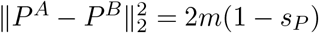 on the strict upper triangle.

“Phase-only” here describes thefinal template coordinates and matching rule, not complete independence from magnitudes throughout feature construction. Since complex averaging occurs before phase extraction, the phase of a given edge is

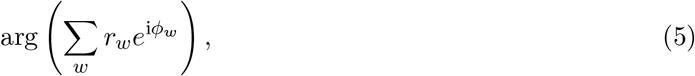

which implicitly weights window phases by their coherency magnitudes. The same issue arises within frequency averaging. An alternative would normalize each edge to unit modulus before averaging, but the supplied retained archive does not establish outcomes for that distinct construction. We therefore do not claim that all amplitude influence has been removed. Phase is undefined at exact zero coherency and can be unstable near zero; the original numerical convention must be documented before release. No new threshold is introduced here.

The normalized real complex inner product on strict-upper entries is

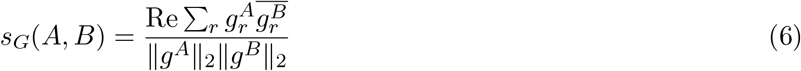

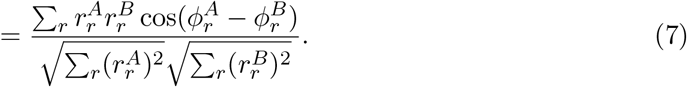

This is a cosine similarity after concatenating real and imaginary coordinates. Its range is [− 1, 1]. The diagonal is excluded from this score, sometimes referred to as “Frobenius cosine.” Using all off-diagonal entries would give the same normalized value by Hermitian symmetry.

The role of magnitudes can be stated more precisely than simply calling them weights. Let 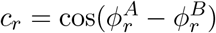 and 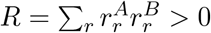. Then

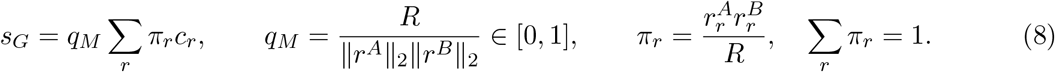

Thus *s*_*G*_ is the product of magnitude-pattern cosine alignment and a magnitude-weighted average of phase agreement. Equivalently, its direct phase weights sum to *q*_*M*_, not generally to one. Edges strong in both recordings receive greater weight, while magnitude-pattern mismatch contracts the score toward zero (which need not reduce a negative score). When all edge magnitudes are equal within each recording, *s*_*G*_ = *s*_*P*_. For nonzero templates with *R* = 0, *s*_*G*_ = 0 although the normalized *π*_*r*_ are undefined. No claim that larger magnitudes are optimal reliability weights follows from this identity.

### 2.4 Holonomy and additional representations

Under node-wise offsets, *ϕ*_*ij*_ 1 ↦*ϕ*_*ij*_ + *θ*_*i*_ − *θ*_*j*_. For every *i < j < k*, define

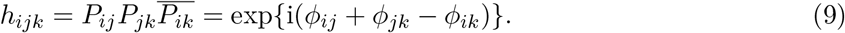

The offset terms cancel exactly. The holonomy score is

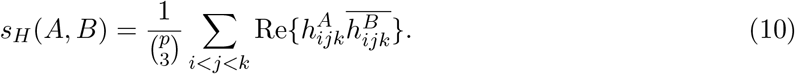

**Figure 1.**
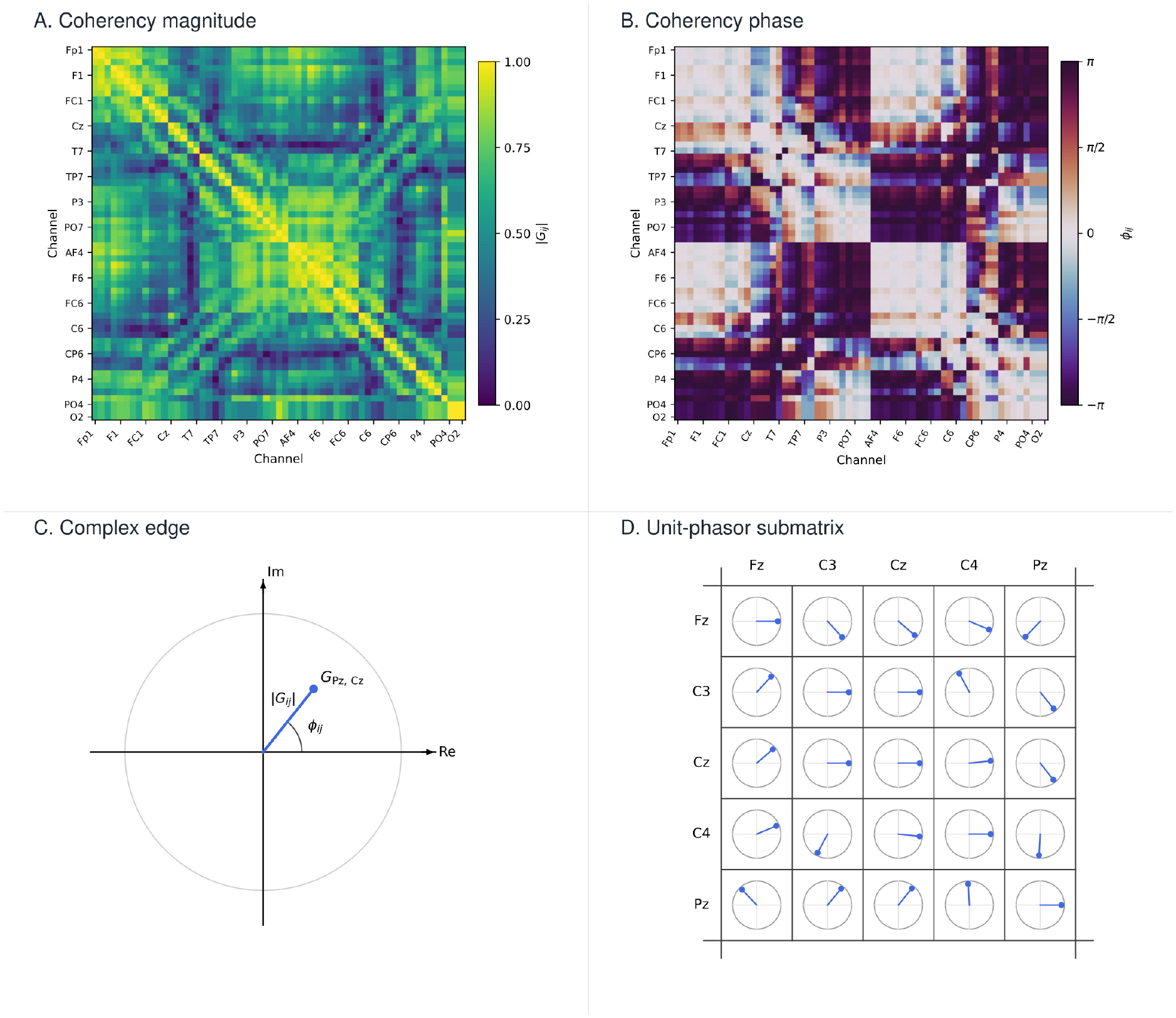
Geometric views of the same archived alpha-band enrollment template (source label: subject 35).**(A)** Coherency magnitude |*G*_*ij*_|. **(B)** Circular phase *ϕ*_*ij*_ = arg *G*_*ij*_. **(C)** Polar form of a single complex entry. **(D)** Five-channel fragment of the unit-phasor matrix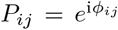. The full-montage channel order is preserved in A and B; D is only a visual fragment, not a reduced channel panel used for identification. The source graphics do not establish the exact visit or recording support. Panels were assembled from the supplied vector figures using AI-assisted layout editing; the numerical source data were not altered.

All 32,509 RestCog triangles, 41,664 Dortmund triangles, or 26,235 L-FAME triangles are included, with no selected reference channel or cycle subset. These coordinates are redundant: the complete graph has *m* − *p* + 1 independent cycle degrees of freedom (1653, 1953, and 1431, respectively). Equality of all triangle holonomies determines the same edge-phase gauge class, provided all phases exist; a proof is included in the Supplementary Material. The mean triangle score is a particular metric-like comparison of this redundant representation, not the result of minimizing edge distance over node phases. Gauge invariance does not imply invariance to general channel mixing or to a change of EEG reference.

Write the independent entries as 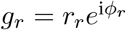, with *r*_*r*_ = | *g*_*r*_|. The primary magnitude comparator in the state panels is negative root-mean-square distance,

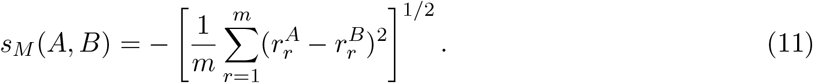

Magnitude Pearson and cosine controls are reported separately where available. Magnitude here is 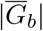, not magnitude-squared coherence and not the average of window magnitudes.

### 2.5 Matching, fusion, and performance metrics

#### 2.5.1 Fixed band and representation fusion

Rows of an *N* × *N* score matrix index probe participants and columns index enrollment participants. Row-wise standardization and equal alpha–beta fusion are

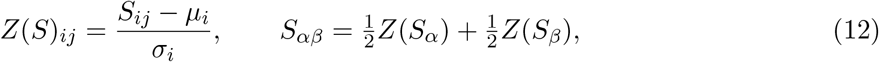

where *µ*_*i*_ and *σ*_*i*_ are computed over the enrollment candidates in probe row *i*. The archive documents fixed 0.5*/*0.5 weights without label-fitted calibration. The exact zero-variance safeguard and standard-deviation convention await code reconciliation. Native holonomy alpha–beta fusion, reported as a secondary control, instead averages the two unstandardized scores. An exploratory theta–alpha–beta fusion was also retained for Dortmund; the corresponding source metrics remain archived but are not used for the principal comparison.

Cross-family fusion uses the same equal row-Z combination of two representation score matrices. For the alpha–beta branch, the band-fused representation scores are subsequently combined at the representation level; the archived script should confirm the exact normalization nesting. A phase-plus-magnitude score fusion is not Eq. (7): it adds normalized scores rather than weighting individual phase edges by pairwise magnitudes.

Row normalization uses no identity labels but does use the complete candidate gallery. For a nonconstant row it preserves its ranking before fusion, while changing the ordering of scores drawn from different probe rows. Therefore it can change pooled verification AUC even for a single representation. Fusion AUC must be interpreted as gallery-normalized verification, not as performance of a gallery-independent pairwise authenticator. Single-band row-Z controls are not available for every condition, so normalization effects and band complementarity cannot always be isolated.

#### 2.5.2 Identification, verification, and descriptive analysis

With 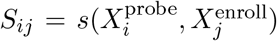, diagonal entries are genuine and off-diagonal entries are impostors. Each probe is ranked independently; there is no global one-to-one assignment constraint. For genuine rank *R*_*i*_,

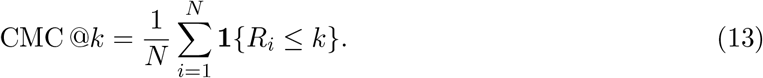

CMC@1 is the fraction correctly identified in a gallery of size *N*. CMC@5 and equal-error rate (EER) are retained when available. Tie handling follows the recorded implementation and requires afinal code-to-text check. One correctly identified participant corresponds to 1.67 percentage points in RestCog and approximately 0.49 in Dortmund.

Verification AUC compares the pooled genuine and impostor score distributions. It summarizes the probability that a randomly chosen genuine score exceeds a randomly chosen impostor score, with half-credit for ties. There are 60 genuine and 3540 impostor scores in each RestCog panel, and 206 genuine and 42,230 impostor scores in Dortmund. These are not independent observations: many comparisons share a recording. EER values in the archive use the nearest available ROC operating point and should not be interpreted as calibrated deployment thresholds.

The results are reported descriptively; no confidence intervals or paired significance tests are claimed. Differences are descriptive, and no preregistration is claimed. Fixed scoring does not remove the exploratory character of comparing several representations and partitions. When used descriptively, a correctness union requires true identity labels and is not an implementable matching score.

### 2.6 Use of AI-assisted tools

During the development of this study and preparation of the manuscript, the corresponding author used ChatGPT (OpenAI; GPT-family models accessed through ChatGPT in 2026) as an assistive tool for methodological discussion, software development and debugging, literature discovery and reference checking, manuscript organization, language revision, L^A^TEX preparation, and layout of figures based on existing data. AI-assisted suggestions and outputs were critically reviewed and, where applicable, checked against the original analysis records and cited sources. The authors retain full responsibility for the study design, implementation, analyses, reported results, and scientific interpretations. No synthetic EEG recordings or empirical results were generated using generative AI.

## 3 Results

### 3.1 Longitudinal persistence across three independent cohorts

Same-state matching across separate visits revealed individual information in both the phase-only and full complex representations in all three cohorts (Table 2). These comparisons were conducted within each cohort’s recording protocol; differences in gallery size, montage, and support preclude treating the three intervals as a shared temporal decay series.

**Table 2.**
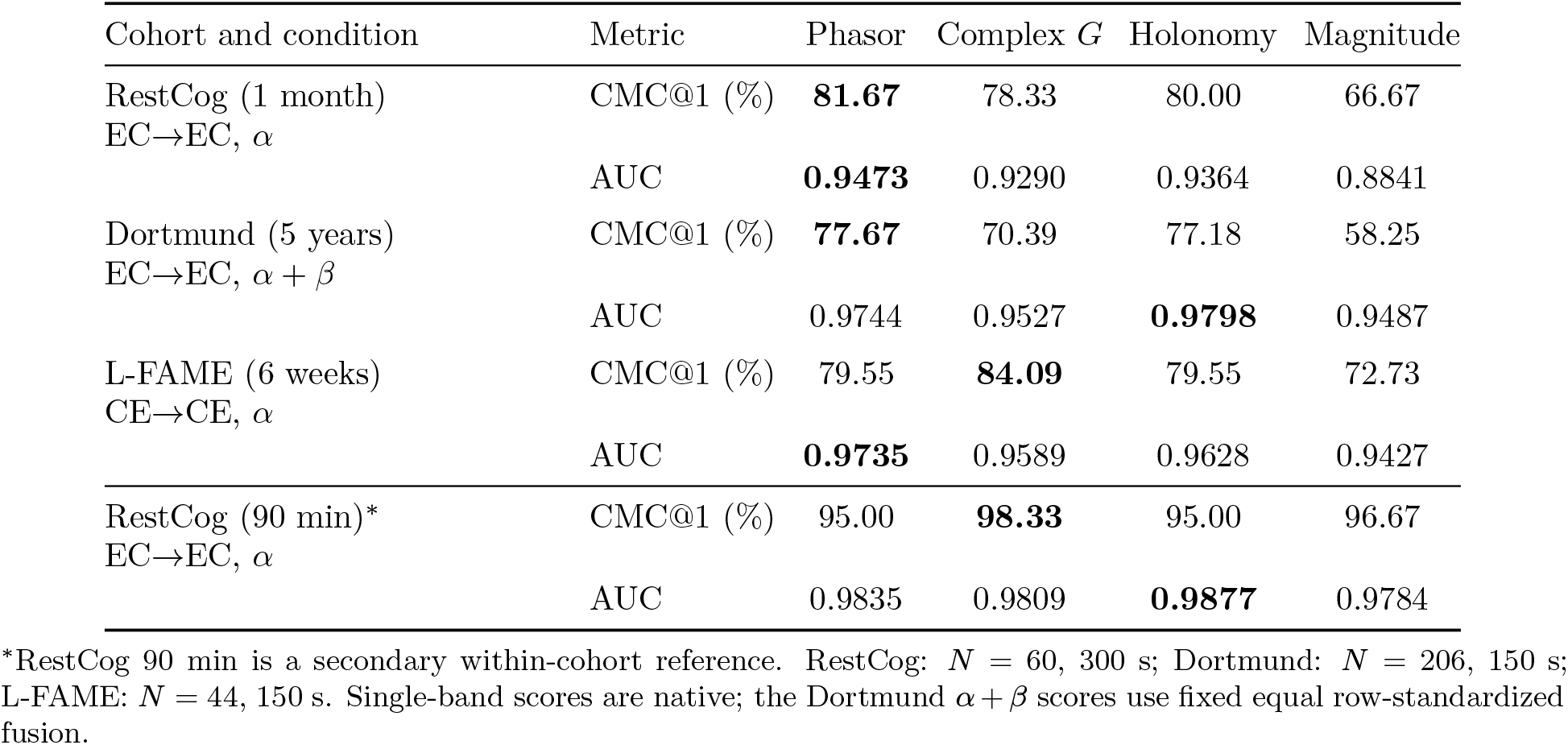
Same-state longitudinal fingerprints. CMC@1 (%) and verification AUC are shown on separate rows; bold indicates the largest value within each row, not a statistically established ranking.

| Cohort and condition | Metric | Phasor | Complex *G* | Holonomy | Magnitude |
|---|---|---|---|---|---|
| RestCog (1 month)<br>EC→EC, *α* | CMC@1 (%) | **81.67** | 78.33 | 80.00 | 66.67 |
|  | AUC | **0.9473** | 0.9290 | 0.9364 | 0.8841 |
| Dortmund (5 years)<br>EC→EC, *α* + *β* | CMC@1 (%) | **77.67** | 70.39 | 77.18 | 58.25 |
|  | AUC | 0.9744 | 0.9527 | **0.9798** | 0.9487 |
| L-FAME (6 weeks)<br>CE→CE, *α* | CMC@1 (%) | 79.55 | **84.09** | 79.55 | 72.73 |
|  | AUC | **0.9735** | 0.9589 | 0.9628 | 0.9427 |
| RestCog (90 min)\*<br>EC→EC, *α* | CMC@1 (%) | 95.00 | **98.33** | 95.00 | 96.67 |
|  | AUC | 0.9835 | 0.9809 | **0.9877** | 0.9784 |

<sup>\*</sup>RestCog 90 min is a secondary within-cohort reference. RestCog: *N* = 60, 300 s; Dortmund: *N* = 206, 150 s; L-FAME: *N* = 44, 150 s. Single-band scores are native; the Dortmund *α* + *β* scores use fixed equal row-standardized fusion.

The representation ranking was not uniform. In monthly RestCog, alpha phasors gave the highest rank-1 identification and AUC, closely followed by holonomy and full complex coherency. In L-FAME, alpha complex coherency identified more individuals than phasors, while phasors provided better verification separation. The longest interval, approximately five years in Dortmund, yielded 77.67% rank-1 identification with fixed alpha–beta phasor fusion; holonomy gave a slightly higher AUC under the same protocol. Magnitude-only matching was less effective in these principal longitudinal comparisons.

RestCog also provides a short-interval reference: identification at approximately 90 minutes was near ceiling for all four representations, but lower at one month. The later decline was not identical across representations. Because the visits may also differ in acquisition and physiology, this within-cohort contrast is informative about persistence without isolating a rate of temporal decay.

An independent Dortmund Pearson baseline yielded lower CMC@1 and AUC than alpha phasors (Supplementary Table S1), although preprocessing differed.

### 3.2 Functional-state dependence of longitudinal identification

Changing the functional state challenged the longitudinal fingerprints in different ways. In RestCog, eyes-closed enrollment was kept fixed while the month-separated probe shifted from rest to episodic recall (ME), internally generated music (MU), or mental arithmetic (MA) (Figure 2). Alpha phasor identification decreased most for arithmetic, whereas full complex coherency showed smaller rank-1 losses relative to its own rest baseline in the MU and MA comparisons. Phasors nevertheless retained the higher verification AUC in every monthly probe state. These results point to different state sensitivities of rank-1 matching and verification, rather than a uniformly superior representation.

**Figure 2.**
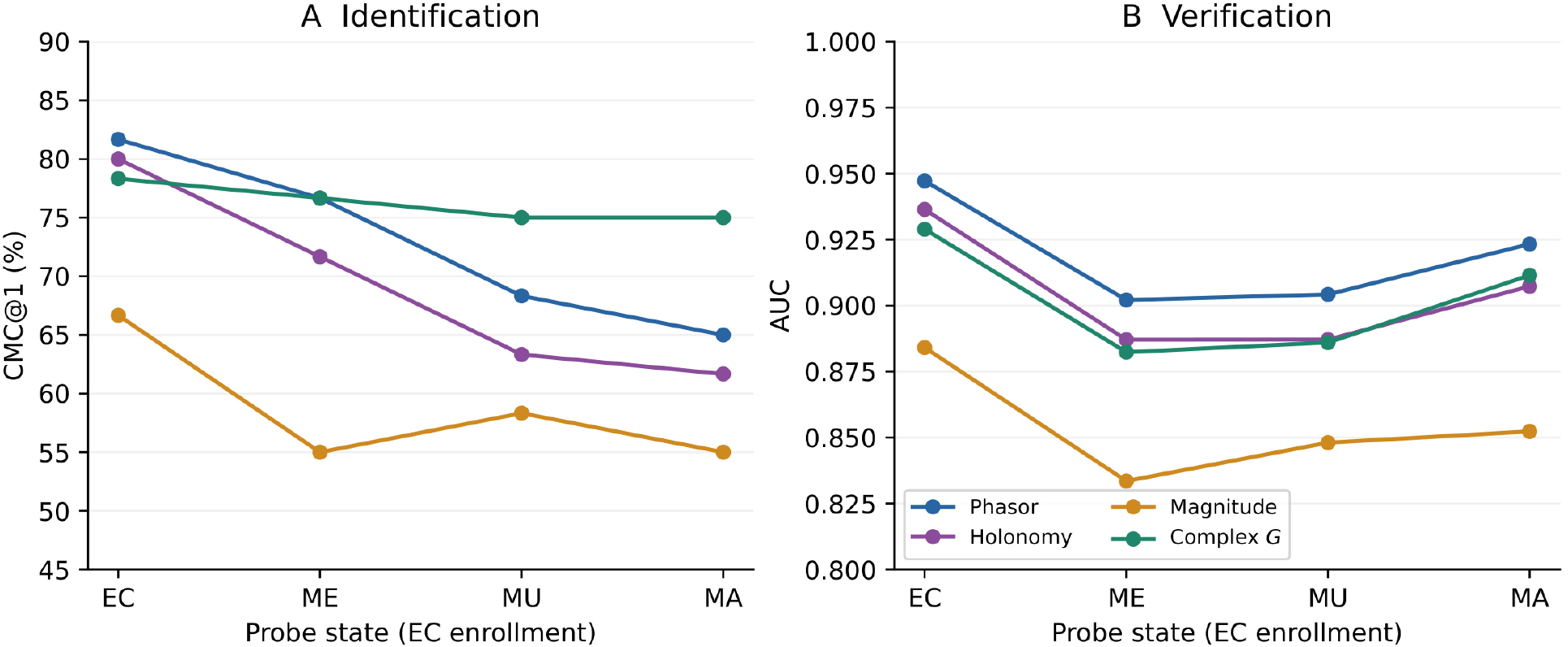
Monthly RestCog EC enrollment: alpha-band identification (A) and verification (B) across probe states. All points use *N* = 60 and 6 × 50 s. Lines connect categorical conditions, not successive times; no participant-level uncertainty estimates are implied.

A more pronounced state-transfer penalty appeared in Dortmund when participants changed between eyes-closed (EC) and eyes-open (EO) recordings (Table 3). Same-state matching remained substantially more accurate than either cross-state direction for both phasors and full complex coherency. Verification scores retained considerable genuine–impostor separation despite the much weaker rank-1 identification. Thus, persistence over five years does not guarantee invariance to eye condition. The available protocol does not include a short-interval EC/EO comparison that could separate a state-by-time interaction.

**Table 3.** Dortmundfive-year EC/EO transfer at 150 s, *α* + *β* equal row-Z fusion.

| Enroll→probe | Phasor | Holonomy | Magnitude | Complex *G* |
|---|---|---|---|---|
| *Identification: CMC@1 (%)* |  |  |  |  |
| EC→EC | 77.67 | 77.18 | 58.25 | 70.39 |
| EC→EO | 29.61 | 30.10 | 14.56 | 23.79 |
| EO→EC | 22.33 | 21.84 | 16.02 | 16.99 |
| EO→EO | 61.17 | 61.65 | 50.00 | 54.85 |
| *Verification: AUC* |  |  |  |  |
| EC→EC | 0.9744 | 0.9798 | 0.9487 | 0.9527 |
| EC→EO | 0.8291 | 0.8504 | 0.7596 | 0.8009 |
| EO→EC | 0.8270 | 0.8337 | 0.7612 | 0.7746 |
| EO→EO | 0.9315 | 0.9343 | 0.9044 | 0.9018 |

*N* = 206, 64 channels. CMC@5 and EER are not available for this new state panel. Enrollment is visit 1 and probe is visit 2 in every direction.

L-FAME instead compares two eyes-closed conditions, rest and silent meditation, across the six-week visit interval (Figure 3). Cross-state identification remained comparatively strong. With fixed alpha–beta fusion, full complex coherency reached 86.36% rank-1 identification for rest-to-meditation transfer and also performed well for meditation-to-meditation matching. Phasors and holonomy were similarly informative in other cells. The different behavior of the two cohorts should not be attributed to eye opening alone: their intervals, acquisition settings, and participant groups also differ.

**Figure 3.**
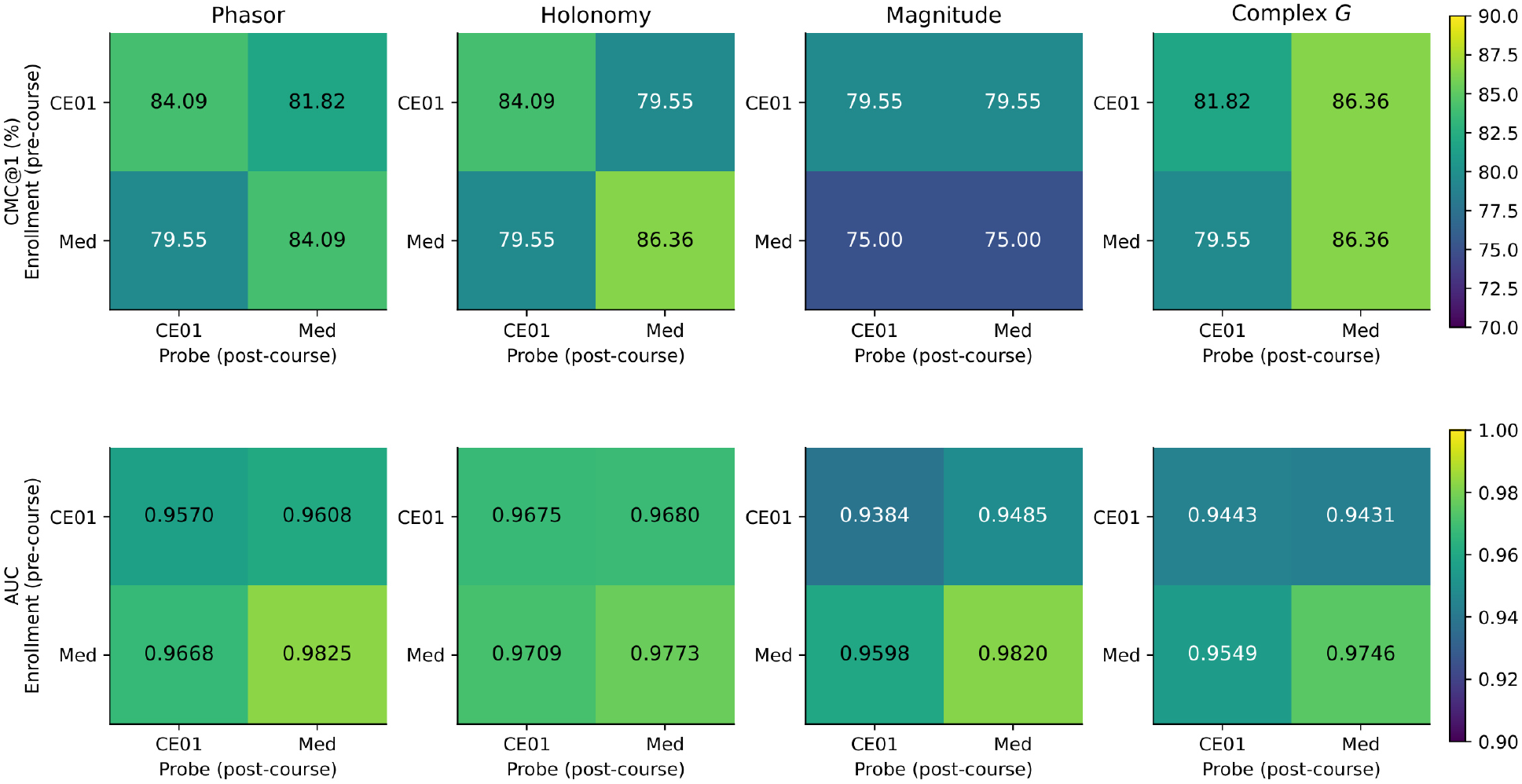
Six-week L-FAME state transfer at 150 s using fixed alpha–beta fusion. Rows denote pre-course enrollment and columns post-course probes; both directions preserve chronological visit order. The top panels show CMC@1 (%), the bottom panels pooled AUC. CE01 denotes eyes-closed rest and Med silent meditation; colors encode aggregate scores, not subject-level similarity matrices.

### 3.3 Complementary information in phase, complex coherency, and holonomy

The varying rankings can be understood in light of the geometry of the two central scores. Phasor matching compares edge-phase differences with equal weights (Eq. (4)); full complex cosine also accounts for the agreement of magnitude patterns and weights each phase comparison by the product of its edge magnitudes (Eq. (8)). In the monthly RestCog arithmetic comparison, the complex score ranked the true identity first more often, although phasors achieved better pooled AUC. The same split between identification and verification appeared in L-FAME rest-to-rest matching. Dortmund, by contrast, favored phasors over complex coherency on both criteria under alpha–beta fusion. The main tables and figures therefore support complementary, not hierarchical, descriptions of the individual fingerprint.

Holonomy preserved substantial individual information after factoring out node-wise phase off-sets. Its performance near phasors in the same-state comparisons shows that the fingerprint is not confined to phases removable by these offsets. This algebraic invariance, however, neither removes volume conduction nor renders the sensor-level descriptor EEG-reference independent. Magnitude alone was less discriminative in the primary RestCog and Dortmund comparisons, but its effect within the complex score cannot be inferred from magnitude-only matching. The supplementary additive-fusion controls likewise show that combining two scores can improve pooled AUC while reducing rank-1 identification.

CMC@1 measures whether the genuine identity wins within a particular gallery row, whereas pooled AUC measures separation across genuine and impostor comparisons from all rows. Disagreement between the two is therefore meaningful, rather than an inconsistency. Differences among representations are descriptive; participant-level paired uncertainty was not estimated.

### 3.4 Frequency fusion and recording support

Alpha phasors formed the strongest single-band fingerprint in the reported comparisons. Beta added useful complementary ranking information through fixed alpha–beta fusion in Dortmund and L-FAME, but the same fusion sometimes reduced monthly RestCog rank-1 identification. Row standardization is part of that fusion procedure and also affects pooled AUC, so a gain in verification cannot be assigned to beta alone without matched normalized single-band controls.

Longer recording support generally strengthened identification, although its effect depended on the representation (Figure 4). In L-FAME, alpha-phasor identification improved from 50 to 100 to 150 s in each of the four directed state pairs. Several fused trajectories were non-monotonic, illustrating that adding data does not necessarily improve a fixed combined score. Monthly RestCog likewise showed better alpha-phasor identification at 300 s than at 150 s; its three 300-s partitions agreed in aggregate rank-1 accuracy even though the partitions use different spectral smoothing bandwidths. The detailed duration and fusion results are retained in the Supplementary Material.

**Figure 4.**
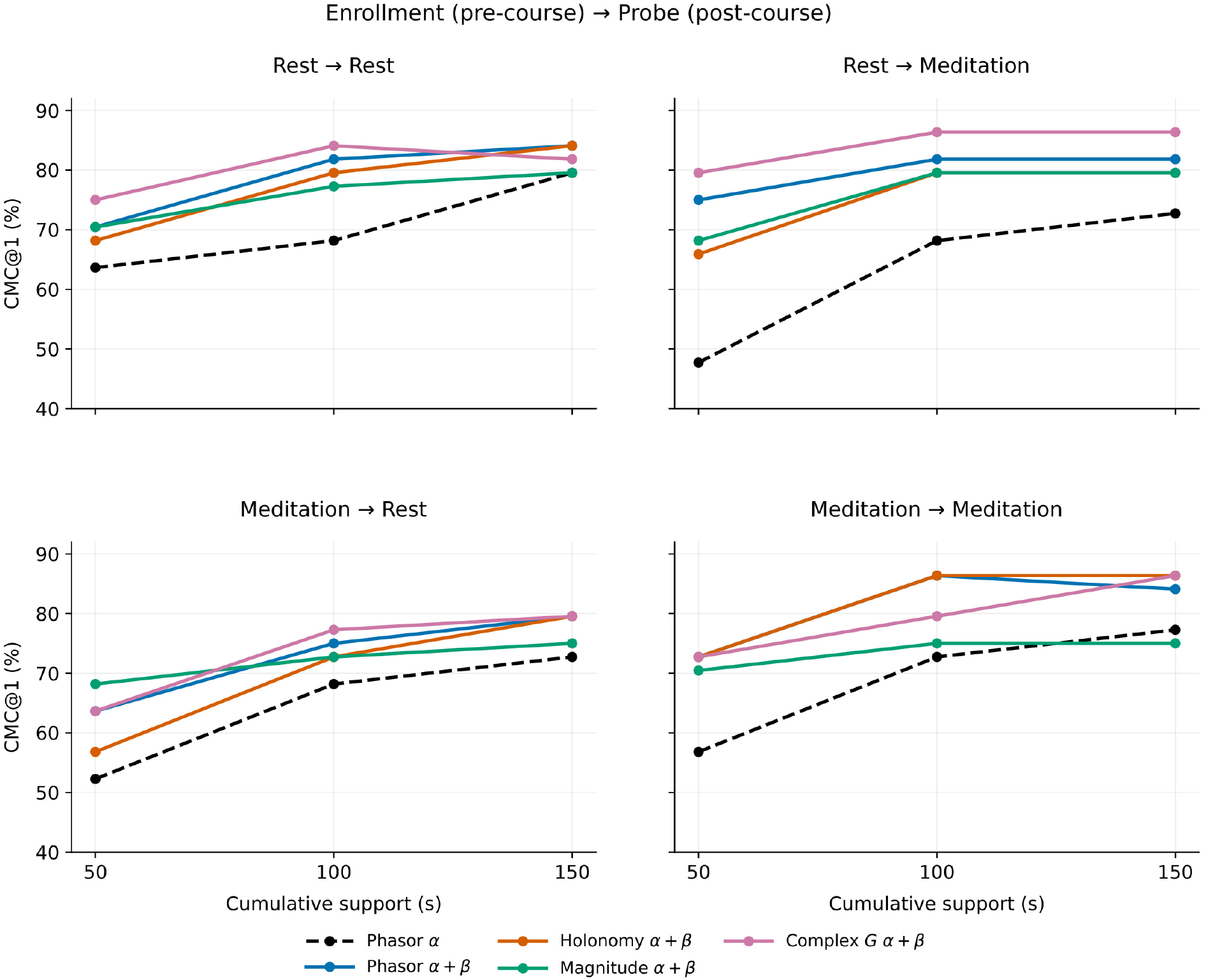
L-FAME rank-1 identification versus cumulative recording support (50, 100, and 150 s). Panel titles give enrollment state (pre-course) → probe state (post-course): Rest = CE01 (eyes-closed rest), Meditation = slMedita (silent meditation). The black dashed line denotes alpha phasors; solid lines denote fixed alpha–beta fusion of phasors, holonomy, magnitude, and full complex coherency. Supports are nested recordings, not independent samples. Panel labels and spacing were adjusted with AI-assisted plotting code; the archived values were unchanged.

## 4 Discussion

### 4.1 Individual EEG geometry persists across visits

An EEG fingerprint need not be identical at two visits to remain identifiable. Across three independent cohorts, inter-channel coherency phases retained sufficient individual structure for direct matching after intervals ranging from weeks to approximately five years. The full complex-coherency representation also remained informative, sometimes matching more individuals than phasors and sometimes fewer. Neither representation required a learned identity classifier.

Recent EEG biometric approaches place complementary emphasis on learned features or spatial graph structure. Li et al. [13] used electrode geometry in connectivity-network individualization, Tian et al. [20] fused multiband PLV-derived graphs using a graph convolutional network, and Oztemel and Soysal [15] investigated attention-autoencoder features and downstream classifiers over two visits. Their accuracy values are not directly comparable with ours because their datasets, gallery sizes, learning protocols, and recording conditions differ. The distinction of the present study is a fixed, directly interpretable comparison of signed coherency phase, full complex coherency, holonomy, and magnitude over substantially different longitudinal intervals and functional states.

The strongest evidence comes from comparisons between separate visits rather than repeated recordings within one session. In RestCog, alpha phasors identified 49 of 60 participants after approximately one month, compared with 47 of 60 for full complex coherency. In L-FAME, the corresponding six-week rest-to-rest counts were 35 and 37 of 44. Dortmund extends the observation to a much longer interval: with fixed alpha–beta fusion, phasors identified 160 of 206 participants approximately five years after enrollment. Together, these results show that both phase and complex representations can preserve useful individual information across substantial gaps between recordings.

The 90-minute RestCog comparison helps put the monthly result in perspective. Identification was higher at the shorter interval for both representations, even though the cohort, channel panel, and analyzed support were the same. The difference is consistent with reduced reproducibility across more widely separated visits, but elapsed time cannot be disentangled from other session-specific changes. Likewise, Dortmund, RestCog, and L-FAME differ in participants, acquisition settings, montage sizes, and gallery sizes. Their results establish persistence in distinct longitudinal settings, not a common decay curve.

Related findings from these cohorts help distinguish identity matching from other forms of longitudinal stability. In RestCog, the reliability and subject distinctiveness of frozen EEG embeddings were not interchangeable properties [7]. Five-year intraclass correlations of spectral markers in Dortmund establish repeatability of those measures, rather than correct identity assignment in a gallery [16]. Likewise, L-FAME’s published classification and cross-session adaptation benchmarks do not evaluate person recognition [12]. Our rank-based and verification outcomes complement these studies; differences in targets and protocols preclude direct numerical comparisons.

A separately evaluated correlation-matrix baseline on Dortmund provides a conventional-connectivity reference, but preprocessing differences prevent a strictly controlled phase-versus-correlation inference (Supplementary Table S1).

The measurement-level character of this result matters. Stable head anatomy, electrode-placement conventions, referencing, and persistent non-neural contributions may all support person recognition alongside neural activity. Our findings extend work on EEG connectivity-based identification [6, 21] by separating the roles of signed phase and complex magnitude in matching, not by identifying a unique cortical mechanism of individual differences.

### 4.2 Functional state changes the fingerprint, but not in a uniform way

A phase pattern that persists across visits may still respond to the participant’s functional state. This distinction is apparent in RestCog, where enrollment remains eyes-closed rest while the monthly probe changes to memory recall, internally generated music, or mental arithmetic. For the latter condition, alpha-phasor rank-1 identification declined from 81.67% in rest to 65.00%, whereas full complex coherency changed from 78.33% to 75.00%. The complex score showed smaller losses relative to its own rest baseline in several cognitive-state comparisons. Yet phasors retained higher verification AUC across all four monthly probe conditions, including mental arithmetic (0.9233 versus 0.9115). Thus the relative advantage depends not only on state but also on what the evaluation measure rewards.

The two additional cohorts put these results in context. In Dortmund, switching between eyes-closed and eyes-open recording caused a much larger reduction in identification than matching the same eye condition, for phasors and complex coherency alike. L-FAME showed comparatively strong transfer between eyes-closed rest and silent meditation; in the rest-to-meditation direction, alpha– beta complex fusion reached 86.36% CMC@1, against 81.82% for phasor fusion. Opening the eyes and changing internally directed mental activity are clearly different manipulations, so their effects should not be treated as equivalent tests of state invariance.

Several boundaries to this interpretation follow directly from the designs. Monthly RestCog enrollment is available only in EC, and the task conditions differ in cognitive content [22]. L-FAME compares visits before and after a meditation course, without isolating training effects from other between-visit changes. Nor can the different cohorts establish whether the effect of state depends on elapsed time. What the data show is more specific: longitudinal identity information survives several state transitions, but its expression depends on the recording condition and on the geometry used for matching.

### 4.3 What phase, magnitude, and holonomy contribute

The contrast between phasor and complex-coherency matching has a precise mathematical basis. The phasor score compares edge-phase differences with equal weight at the final matching stage. By contrast, Eq. (8) expresses complex cosine as magnitude-pattern alignment multiplied by a magnitude-weighted average of phase agreement. Edges with large coherency magnitudes in both recordings contribute more to the complex score, while disagreement between magnitude patterns changes its overall scale. These are different geometries of the same complex matrix, not inter-changeable ways of implementing a phase metric.

This decomposition helps interpret the mixed empirical ordering. Downweighting weak edges could reduce the influence of poorly determined phases and may help explain why complex coherency maintained higher rank-1 identification in some RestCog state transfers. But coherency magnitude is not a validated measure of edge reliability: strong values can arise through mixing or common sources, and weak edges may still be individually distinctive. The aggregate results cannot determine which edges account for the observed differences. Magnitude-only matching also performed less well than the principal phase and complex scores in RestCog and Dortmund, showing that the contribution of magnitude within *s*_*G*_ cannot be inferred from magnitude-only performance.

An important qualification applies even to the term “phase-only.” Phase is extracted after complex frequency and window averaging, so the estimated phase already reflects the magnitudes of the terms being averaged. Phasor and complex matching therefore differ in their final comparison rule; the analysis does not isolate phase from every amplitude-dependent operation earlier in the pipeline. Likewise, additive fusion of separately normalized phase and magnitude scores does not reproduce the multiplicative structure of the complex cosine, and the available fusion results show no uniform gain.

Holonomy supplies a complementary geometric perspective. Its triangle phases are unchanged by node-wise phase offsets, and their successful longitudinal matching indicates that individual information survives this algebraic reduction. If the edge phases consisted only of differences between node potentials, every triangle holonomy would be one; the observed discrimination therefore reflects more than such a pure-offset pattern. This is an invariance of the representation, not evidence of directed physiological circulation. In particular, triangle holonomy is neither invariant to arbitrary scalp-signal mixing nor a remedy for volume conduction or reference effects [2]. The many overlapping triangles also form a redundant set of coordinates rather than independent physiological observations.

### 4.4 Frequency fusion and evaluation criteria

Alpha carried the strongest single-band phasor fingerprint in the reported RestCog, Dortmund EC, and L-FAME comparisons. Beta nevertheless provided complementary information in some fixed alpha–beta fusions, notably in Dortmund and L-FAME. RestCog is a useful counterexample to any general fusion rule: monthly phasor CMC@1 fell from 81.67% for raw alpha to 75.00% after row-standardized alpha–beta fusion. Extra frequency information can improve matching, but its benefit depends on the cohort, condition, and score construction.

Identification and verification also test different aspects of a score matrix. CMC@1 asks whether the true enrollment recording ranks above every competing identity for a given probe. Pooled AUC instead compares genuine scores with impostor scores across the gallery. One difficult competitor can change a participant’s rank-1 outcome without producing an equivalent change in pooled score separation. This explains why phasors and full complex coherency sometimes reverse order between CMC@1 and AUC, rather than revealing an inconsistency in the results.

Row-wise standardization adds a further distinction. It preserves rankings within each single-band probe row, but changes score location and scale across probe rows before fusion. As a result, an increase in pooled AUC after alpha–beta fusion cannot be assigned entirely to the beta band without corresponding row-standardized single-band controls. These gallery-dependent scores are appropriate for the closed-gallery comparisons studied here; they have not been validated for standalone authentication, open-set recognition, or a fixed deployment threshold. High verification AUC in this setting should not be read as a biometric security guarantee.

### 4.5 Scope and limitations

Several features of the design define the strength and the limits of these conclusions. The three cohorts differ in recording support, montage, age composition, and gallery size. RestCog provides a shorter within-cohort reference and monthly EC-to-task comparisons, Dortmund offers five-year EC/EO comparisons, and L-FAME provides rest/meditation transfer at nested recording durations. These protocols complement one another but do not separate time, state, and acquisition effects in a common factorial design. The RestCog monthly panel lacks task-state enrollment, and its EO analysis was not completed. Some additional state-transfer panels also lack rank-5 or equal-error summaries, narrowing the comparisons available for those outcomes.

The analysis is descriptive. No participant-level paired significance tests or uncertainty intervals were calculated for the representation contrasts, and the many genuine–impostor comparisons share the same recordings. Treating those scores as independent observations would overstate precision. Comparisons among representations, bands, and fusion schemes should be viewed as exploratory rather than preregistered confirmatory tests, despite fixed matching rules and the absence of a learned identity classifier. The duration analyses are likewise nested rather than independent, and changing the RestCog window partition also changes multitaper spectral smoothing.

Finally, interpretation remains bounded by the scalp recording model. Volume conduction, common input, reference choice, and recording noise can influence the observed phase and magnitude relationships [2, 14, 18]. Phase is also undefined for zero coherency and may be unstable for very small magnitudes. Our algebraic holonomy reduction cannot resolve these measurement effects, and no source-reconstruction or anatomy-controlled analysis was performed. The results therefore quantify persistent person-specific discriminability in measured EEG, not the fraction attributable to stable cortical interactions. Within that scope, phase-only and full complex geometry retain complementary information, with different responses to state changes and evaluation criteria.

## 5. Conclusion

Across three independent EEG cohorts, inter-channel phase patterns and full complex coherency supported direct identification without a trained classifier, with visits separated by weeks to approximately five years. Persistence was nevertheless sensitive to functional state. In Dortmund, transfer between eyes-closed and eyes-open recordings sharply reduced identification, whereas RestCog and L-FAME retained substantial individual information across changes in cognitive condition.

Unit phasors and full complex coherency offered complementary descriptions of this persistence. Phasor matching emphasizes relative-phase agreement, while the full complex score also reflects magnitude-pattern alignment and magnitude-weighted phase agreement. Their relative advantages changed with the recording condition and with the choice between rank-1 identification and verification AUC. Triangle holonomies retained discriminative information after node-wise phase freedom was factored out. The resulting fingerprints are persistent but state-sensitive properties of recorded scalp EEG coherency; their identification performance does not by itself establish a uniquely neural source for the individual differences.

## Acknowledgments

The authors thank the investigators and participants of the RestCog, Dortmund Vital Study, and L-FAME projects for making the longitudinal EEG datasets available for research. The corresponding author used ChatGPT (OpenAI; GPT-family models accessed via ChatGPT in 2026) for methodological discussion, coding assistance, literature and reference checks, manuscript organization, language editing, and arranging existing figures. AI-assisted material was critically reviewed by the authors; responsibility for the analyses,findings, and manuscript remains with the authors. The use of AI-assisted tools is also described in Materials and Methods.

## Statements and Declarations

### Author contributions

D.G.: Conceptualization, Methodology, Investigation, Formal analysis, Data curation, Writing – original draft. A.D., B.N., and S.S.: Investigation, Data curation, Validation, Visualization, Formal analysis. N.J.Z.: Conceptualization, Methodology, Supervision, Project administration, Writing – review and editing.

### Ethics approval

This study is a secondary analysis of previously collected, publicly available EEG datasets; no new human recordings were acquired. The original RestCog study reported ethics approval and informed consent [22]. The Dortmund Vital Study reported approval by the local ethics committee at IfADo (A93-1 and A93-3) and written informed consent [8]. L-FAME reports approval by the Michigan State University Institutional Review Board and written informed consent [10].

### Consent to participate

Participant consent was obtained in the original studies, as described in their respective dataset documentation. No additional participants were recruited for this secondary analysis.

### Data availability

The EEG datasets analyzed in this study are publicly available: RestCog through OpenNeuro (https://openneuro.org/datasets/ds004148), the Dortmund Vital Study through OpenNeuro (https://openneuro.org/datasets/ds005385), and L-FAME through its dataset repository (https://huggingface.co/datasets/L-FAME-Dataset-Benchmark/L-FAME). The numerical results are summarized in the article and the accompanying Supplementary Information. The manuscript source archive contains additional aggregate tables and numerical outputs used in preparing the figures and tables.

### Code availability

Scripts for regenerating manuscript tables and figures from the accompanying aggregate data are included in the source archive. The EEG preprocessing and feature-extraction code will be made available to editors and reviewers during peer review. A curated release with software dependencies and reproducibility instructions is planned for deposition in a public repository upon acceptance.

