## Supplementary Information for "Longitudinal EEG identification using sensor-level phase and complex coherency across sessions and functional states"

This file contains selected mathematical details and extended data tables supporting the primary manuscript. Descriptive values are transcribed from the retained experimental archive. Full machine-readable aggregate results and output provenance are retained separately; absent metrics are not inferred. No additional EEG analysis was performed to prepare this supplement.

### S1. Mathematical properties of the coherency scores

#### S1.1. Triangle holonomies and gauge equivalence

Let $P$ and $Q$ be complete Hermitian unit-phasor edge assignments with identical triangle holonomies and define $R_{ij} = Q_{ij}\overline{P_{ij}}$. Equality of holonomies implies $R_{ij}R_{jk}R_{ki} = 1$ for every triangle. Fix node 1 and put $u_i = R_{i1}$, $u_1 = 1$. On triangle $(i, 1, j)$, $R_{ij} = R_{i1}R_{1j} = u_i\overline{u_j}$, so $Q_{ij} = u_iP_{ij}\overline{u_j}$. Conversely, node-wise phase rotations cancel around every triangle. A common phase of all $u_i$ is immaterial, leaving $p - 1$ gauge degrees of freedom and $m - p + 1$ cycle degrees of freedom. This algebraic equivalence is not a claim about physiological sources or reference invariance.

#### S1.2. Complex distance decomposition

For each edge $g_r = r_re^{i\phi_r}$, the exact identity

$$|g_r^A - g_r^B|^2 = (r_r^A - r_r^B)^2 + 2r_r^Ar_r^B[1 - \cos(\phi_r^A - \phi_r^B)]$$

separates magnitude difference from magnitude-weighted phase difference. Negative complex RMS is distinct from normalized complex cosine and is not used as its substitute.

#### S2. Independent Pearson-correlation baseline: Dortmund

For the same nominal Dortmund study design (session 1 EC enrollment, session 2 EC probe, $N = 206$, 64 channels, 150 s per recording), we retained an independently completed conventional correlation-matrix analysis. Recordings were resampled from 1000 to 128 Hz and alpha-band filtered at 8–13 Hz using a fourth-order Butterworth filter applied with `sosfiltfilt`. Three non-overlapping 50-s windows were used, taking the first three valid windows of each recording. The session template was the arithmetic mean of the three window-wise Pearson correlation matrices, not a spectral coherency matrix. No subject classifier, learned fusion, rank scan, or score calibration was used.

Three fixed readouts compared probe to enrollment templates: Pearson similarity between strict-upper-triangle vectors, full-matrix Frobenius distance, and affine-invariant Riemannian distance (AIRM). For AIRM, matrices were regularized with $R_{\text{spd}} = 0.99\bar{R} + 0.01I$ before distance calculation. Higher edge-Pearson scores indicate stronger matches; lower distances indicate stronger matches for the two distance metrics. The pooled verification AUC was calculated after negating distances.

The control configuration reports two pre-metric exclusions: `sub-230` (session-1 recording shorter than 150 s) and `sub-425` (severe signal saturation). Its 206 enrollment and probe IDs, score/distance matrices, and operator cache are retained with the project archive. The primary coherency manuscript reports the same cohort size and channel count, but its published summary does not establish an exact subject-ID intersection or identical window locations. In addition, the Pearson pipeline uses a separate alpha-band filter instead of the principal frequency-wise multitaper construction. This comparison is descriptive and does not attribute the difference uniquely to phase information.

**Supplementary Table S1:** Independent Pearson correlation-matrix baseline for five-year Dortmund EC→EC identification ($N = 206$, 64 channels, $3 \times 50$ s per visit; alpha 8–13 Hz). All metrics were independently recomputed from the retained $206 \times 206$ score or distance matrices. This separately processed time-domain control is not a matched-preprocessing ablation against the multitaper phasor method.

| Correlation-matrix matcher | CMC@1 (%) | CMC@5 (%) | AUC | Mean rank |
|---|---|---|---|---|
| Edge-vector Pearson similarity | 47.09 | 61.65 | 0.8994 | 16.85 |
| Frobenius distance | 41.75 | 66.99 | 0.9050 | 15.22 |
| Affine-invariant Riemannian distance | 28.16 | 42.23 | 0.8110 | 30.43 |

#### S3. Full longitudinal and state-transfer tables

**Supplementary Table S2:** Five-year Dortmund EC matching: $N = 206$, 64 channels, $3 \times 50$ s (150 s).

| Representation | CMC@1 (%) | CMC@5 (%) | AUC | EER (%) |
|---|---|---|---|---|
| Phasor $\alpha$ | 62.62 | 78.64 | 0.9523 | 10.74 |
| Phasor $\beta$ | 56.31 | 71.84 | 0.8788 | 18.38 |
| Phasor $\alpha + \beta$ | 77.67 | 90.29 | 0.9744 | 7.03 |
| Holonomy $\alpha$ | 62.14 | 77.67 | 0.9409 | 12.84 |
| Holonomy $\beta$ | 56.31 | 74.27 | 0.8906 | 16.92 |
| Holonomy $\alpha + \beta$ | 77.18 | 89.81 | 0.9798 | 5.69 |
| Magnitude $\alpha$ | 51.94 | – | – | – |
| Magnitude $\alpha + \beta$ | 58.25 | – | 0.9487 | – |
| Complex $G$ $\alpha + \beta^{\dagger}$ | 70.39 | – | 0.9527 | – |

Dashes indicate unreported metrics, not zero. †: complex-$G$ CMC@1 is retained from an exact-summary archive entry and requires primary-CSV reconciliation. Magnitude CMC@1 is recoverable from the fusion logs. Magnitude and complex fusion AUCs are newly recovered exact-recap values; CMC@5 and EER for the phase families retain their earlier EC-log provenance. No Dortmund 300-s result is included.

**Supplementary Table S3:** One-month RestCog EC enrollment to each probe state: CMC@1 (%).

| Representation | EC | ME | MU | MA |
|---|---|---|---|---|
| Phasor $\alpha$ | 81.67 | 76.67 | 68.33 | 65.00 |
| Holonomy $\alpha$ | 80.00 | 71.67 | 63.33 | 61.67 |
| Magnitude $\alpha$ | 66.67 | 55.00 | 58.33 | 55.00 |
| Complex $G$ $\alpha$ | 78.33 | 76.67 | 75.00 | 75.00 |
| Phasor $\alpha + \beta$ | 75.00 | 63.33 | 71.67 | 65.00 |
| Holonomy $\alpha + \beta$ | 78.33 | 63.33 | 70.00 | 65.00 |
| Magnitude $\alpha + \beta$ | 58.33 | 48.33 | 51.67 | 56.67 |
| Complex $G$ $\alpha + \beta$ | 68.33 | 65.00 | 63.33 | 65.00 |

All columns use $N = 60$, 59 channels and $6 \times 50$ s. Cognitive-state columns are EC-to-state tests, not monthly same-task tests.

**Supplementary Table S4:** One-month RestCog EC enrollment to each probe state: verification AUC.

| Representation | EC | ME | MU | MA |
|---|---|---|---|---|
| Phasor $\alpha$ | 0.9473 | 0.9021 | 0.9042 | 0.9233 |
| Holonomy $\alpha$ | 0.9364 | 0.8871 | 0.8871 | 0.9073 |
| Magnitude $\alpha$ | 0.8841 | 0.8335 | 0.8481 | 0.8523 |
| Complex $G$ $\alpha$ | 0.9290 | 0.8824 | 0.8860 | 0.9115 |
| Phasor $\alpha + \beta$ | 0.9832 | 0.9506 | 0.9516 | 0.9703 |
| Holonomy $\alpha + \beta$ | 0.9839 | 0.9413 | 0.9513 | 0.9708 |
| Magnitude $\alpha + \beta$ | 0.9347 | 0.8921 | 0.9279 | 0.9117 |
| Complex $G$ $\alpha + \beta$ | 0.9652 | 0.9354 | 0.9415 | 0.9555 |

All columns use $N = 60$, 59 channels and $6 \times 50$ s. Cognitive-state columns are EC-to-state tests, not monthly same-task tests.

**Supplementary Table S5:** L-FAME at 150 s: identification and verification.

| Representation | CE→CE | CE→Med | Med→CE | Med→Med |
|---|---|---|---|---|
| *CMC@1 (%)* | | | | |
| Phasor $\alpha$ | 79.55 | 72.73 | 72.73 | 77.27 |
| Holonomy $\alpha$ | 79.55 | 65.91 | 72.73 | 75.00 |
| Magnitude $\alpha$ | 72.73 | 59.09 | 68.18 | 68.18 |
| Complex $G$ $\alpha$ | 84.09 | 70.45 | 72.73 | 77.27 |
| Phasor $\beta$ | 68.18 | 56.82 | 56.82 | 68.18 |
| Holonomy $\beta$ | 68.18 | 59.09 | 59.09 | 68.18 |
| Magnitude $\beta$ | 59.09 | 65.91 | 59.09 | 70.45 |
| Complex $G$ $\beta$ | 61.36 | 61.36 | 61.36 | 70.45 |
| Phasor $\alpha + \beta$ | 84.09 | 81.82 | 79.55 | 84.09 |
| Holonomy $\alpha + \beta$ | 84.09 | 79.55 | 79.55 | 86.36 |
| Magnitude $\alpha + \beta$ | 79.55 | 79.55 | 75.00 | 75.00 |
| Complex $G$ $\alpha + \beta$ | 81.82 | 86.36 | 79.55 | 86.36 |
| *AUC* | | | | |
| Phasor $\alpha$ | 0.9735 | 0.9440 | 0.9515 | 0.9650 |
| Holonomy $\alpha$ | 0.9628 | 0.9356 | 0.9382 | 0.9594 |
| Magnitude $\alpha$ | 0.9427 | 0.9142 | 0.9369 | 0.9394 |
| Complex $G$ $\alpha$ | 0.9589 | 0.9309 | 0.9389 | 0.9446 |
| Phasor $\beta$ | 0.8200 | 0.7964 | 0.8169 | 0.8391 |
| Holonomy $\beta$ | 0.8239 | 0.8082 | 0.8289 | 0.8531 |
| Magnitude $\beta$ | 0.8154 | 0.8240 | 0.8085 | 0.8432 |
| Complex $G$ $\beta$ | 0.7906 | 0.7805 | 0.7790 | 0.8125 |
| Phasor $\alpha + \beta$ | 0.9570 | 0.9608 | 0.9668 | 0.9825 |
| Holonomy $\alpha + \beta$ | 0.9675 | 0.9680 | 0.9709 | 0.9773 |
| Magnitude $\alpha + \beta$ | 0.9384 | 0.9485 | 0.9598 | 0.9820 |
| Complex $G$ $\alpha + \beta$ | 0.9443 | 0.9431 | 0.9549 | 0.9746 |

$N = 44$, 55 channels; CE = CE01, Med = slMedita. Enrollment is pre-course and probe is post-course (six weeks). Alpha+beta denotes equal row-Z fusion; single bands use raw scores. CMC and EER are percentages, AUC is unitless. All entries are retained console-log metrics.

**Supplementary Table S6:** L-FAME at 150 s: rank-5 identification and equal-error rates.

| Representation | CE→CE | CE→Med | Med→CE | Med→Med |
|---|---|---|---|---|
| *CMC@5 (%)* | | | | |
| Phasor $\alpha$ | 97.73 | 90.91 | 90.91 | 93.18 |
| Holonomy $\alpha$ | 97.73 | 90.91 | 90.91 | 88.64 |
| Magnitude $\alpha$ | 93.18 | 81.82 | 88.64 | 88.64 |
| Complex $G$ $\alpha$ | 97.73 | 88.64 | 95.45 | 90.91 |
| Phasor $\beta$ | 77.27 | 77.27 | 81.82 | 77.27 |
| Holonomy $\beta$ | 77.27 | 77.27 | 79.55 | 81.82 |
| Magnitude $\beta$ | 77.27 | 79.55 | 72.73 | 79.55 |
| Complex $G$ $\beta$ | 84.09 | 77.27 | 79.55 | 77.27 |
| Phasor $\alpha + \beta$ | 93.18 | 90.91 | 93.18 | 95.45 |
| Holonomy $\alpha + \beta$ | 93.18 | 93.18 | 93.18 | 95.45 |
| Magnitude $\alpha + \beta$ | 86.36 | 88.64 | 86.36 | 95.45 |
| Complex $G$ $\alpha + \beta$ | 88.64 | 88.64 | 90.91 | 97.73 |
| *EER (%)* | | | | |
| Phasor $\alpha$ | 7.61 | 11.34 | 13.53 | 9.62 |
| Holonomy $\alpha$ | 7.16 | 14.11 | 13.53 | 8.11 |
| Magnitude $\alpha$ | 11.23 | 15.54 | 15.22 | 16.28 |
| Complex $G$ $\alpha$ | 8.59 | 13.00 | 13.27 | 13.35 |
| Phasor $\beta$ | 20.32 | 24.37 | 24.47 | 26.06 |
| Holonomy $\beta$ | 23.26 | 22.60 | 23.28 | 24.52 |
| Magnitude $\beta$ | 22.91 | 24.84 | 23.76 | 24.34 |
| Complex $G$ $\beta$ | 27.67 | 26.85 | 29.07 | 27.30 |
| Phasor $\alpha + \beta$ | 5.66 | 8.46 | 7.08 | 3.54 |
| Holonomy $\alpha + \beta$ | 6.63 | 9.91 | 6.34 | 3.81 |
| Magnitude $\alpha + \beta$ | 14.06 | 9.28 | 9.20 | 7.11 |
| Complex $G$ $\alpha + \beta$ | 9.41 | 8.38 | 7.35 | 6.63 |

$N = 44$, 55 channels; CE = CE01, Med = slMedita. Enrollment is pre-course and probe is post-course (six weeks). Alpha+beta denotes equal row-Z fusion; single bands use raw scores. CMC and EER are percentages, AUC is unitless. All entries are retained console-log metrics.

### S4. Recording support and fusion controls

**Supplementary Table S7:** RestCog monthly EC matching by temporal support: CMC@1 (%).

| Representation | 150 s: $3 \times 50$ | 300 s: $6 \times 50$ | $3 \times 100$ | $2 \times 150$ |
|---|---|---|---|---|
| Phasor $\alpha$ | 71.67 | 81.67 | 81.67 | 81.67 |
| Phasor $\beta$ | 41.67 | 41.67 | 40.00 | 40.00 |
| Phasor $\alpha + \beta$ | 66.67 | 75.00 | 73.33 | 75.00 |
| Holonomy $\alpha$ | 63.33 | 80.00 | 78.33 | 80.00 |
| Holonomy $\beta$ | 41.67 | 43.33 | 40.00 | 43.33 |
| Holonomy $\alpha + \beta$ | 70.00 | 78.33 | 75.00 | 76.67 |
| Holonomy $\alpha + \beta$ / native | 70.00 | 78.33 | 76.67 | 76.67 |
| Magnitude $\alpha$ | 61.67 | 66.67 | 66.67 | 66.67 |
| Magnitude $\alpha + \beta$ | 63.33 | 58.33 | 58.33 | 63.33 |
| Complex G $\alpha$ | 71.67 | 78.33 | 75.00 | 78.33 |
| Complex G $\alpha + \beta$ | 70.00 | 68.33 | 73.33 | 70.00 |
| Phasor+Holonomy $\alpha$ | 65.00 | 81.67 | 80.00 | 83.33 |
| Phasor+Magnitude $\alpha$ | 70.00 | 76.67 | 80.00 | 78.33 |
| Holonomy+Magnitude $\alpha$ | 70.00 | 76.67 | 80.00 | 78.33 |
| Phasor+Holonomy $\alpha + \beta$ | 70.00 | 78.33 | 73.33 | 76.67 |
| Phasor+Magnitude $\alpha + \beta$ | 70.00 | 75.00 | 75.00 | 78.33 |
| Holonomy+Magnitude $\alpha + \beta$ | 70.00 | 73.33 | 75.00 | 75.00 |

The last three columns hold total duration fixed. The first column uses half as much data and is a separate support condition. Native holonomy fusion averages unstandardized band scores.

**Supplementary Table S8:** L-FAME CMC@1 (%) at nested durations.

| Representation | 50 s | 100 s | 150 s |
|---|---|---|---|
| *CE01→CE01* | | | |
| Phasor $\alpha$ | 63.64 | 68.18 | 79.55 |
| Phasor $\alpha + \beta$ | 70.45 | 81.82 | 84.09 |
| Holonomy $\alpha + \beta$ | 68.18 | 79.55 | 84.09 |
| Magnitude $\alpha + \beta$ | 70.45 | 77.27 | 79.55 |
| Complex $G$ $\alpha + \beta$ | 75.00 | 84.09 | 81.82 |
| *CE01→slMedita* | | | |
| Phasor $\alpha$ | 47.73 | 68.18 | 72.73 |
| Phasor $\alpha + \beta$ | 75.00 | 81.82 | 81.82 |
| Holonomy $\alpha + \beta$ | 65.91 | 79.55 | 79.55 |
| Magnitude $\alpha + \beta$ | 68.18 | 79.55 | 79.55 |
| Complex $G$ $\alpha + \beta$ | 79.55 | 86.36 | 86.36 |
| *slMedita→CE01* | | | |
| Phasor $\alpha$ | 52.27 | 68.18 | 72.73 |
| Phasor $\alpha + \beta$ | 63.64 | 75.00 | 79.55 |
| Holonomy $\alpha + \beta$ | 56.82 | 72.73 | 79.55 |
| Magnitude $\alpha + \beta$ | 68.18 | 72.73 | 75.00 |
| Complex $G$ $\alpha + \beta$ | 63.64 | 77.27 | 79.55 |
| *slMedita→slMedita* | | | |
| Phasor $\alpha$ | 56.82 | 72.73 | 77.27 |
| Phasor $\alpha + \beta$ | 72.73 | 86.36 | 84.09 |
| Holonomy $\alpha + \beta$ | 72.73 | 86.36 | 86.36 |
| Magnitude $\alpha + \beta$ | 70.45 | 75.00 | 75.00 |
| Complex $G$ $\alpha + \beta$ | 72.73 | 79.55 | 86.36 |

All windows are 50 s; the recording template averages the first 1, 2, or 3 complex matrices before extracting phase or holonomy. No row represents an independently sampled duration cohort.

**Supplementary Table S9:** Fixed equal-weight cross-family score fusion for EC matching.

| Cohort/support | Band | Fusion | CMC@1 (%) | AUC | Rescued/lost |
|---|---|---|---|---|---|
| RestCog 300 s | $\alpha$ | $P + M$ | 76.67 | 0.9717 | 1/4 |
| RestCog 300 s | $\alpha$ | $H + M$ | 76.67 | 0.9741 | 1/3 |
| RestCog 300 s | $\alpha$ | $P + H$ | 81.67 | 0.9773 | 0/0 |
| RestCog 300 s | $\alpha + \beta$ | $P + M$ | 75.00 | 0.9777 | 3/3 |
| RestCog 300 s | $\alpha + \beta$ | $H + M$ | 73.33 | 0.9805 | 2/5 |
| RestCog 300 s | $\alpha + \beta$ | $P + H$ | 78.33 | 0.9849 | 2/0 |
| Dortmund 150 s | $\alpha$ | $P + M$ | 64.56 | 0.9670 | 13/9 |
| Dortmund 150 s | $\alpha$ | $H + M$ | 66.02 | 0.9693 | 16/8 |
| Dortmund 150 s | $\alpha$ | $P + H$ | 63.59 | 0.9688 | 4/2 |
| Dortmund 150 s | $\alpha + \beta$ | $P + M$ | 76.21 | 0.9688 | 11/14 |
| Dortmund 150 s | $\alpha + \beta$ | $H + M$ | 77.67 | 0.9735 | 12/11 |
| Dortmund 150 s | $\alpha + \beta$ | $P + H$ | 76.21 | 0.9776 | 3/6 |

Rescued/lost counts are relative to the first named component. They are archived subject counts, not new inferential tests. $P + M$ is score fusion and is not the complex-$G$ score.

#### S5. Secondary RestCog and component comparisons

**Supplementary Table S10:** Short-interval state transfer in RestCog (6 × 50 s): CMC@1 (%).

| Enroll→probe | $P_\alpha$ | $H_\alpha$ | $M_\alpha$ | $G_\alpha$ | $P_{\alpha\beta}$ | $H_{\alpha\beta}$ |
|---|---|---|---|---|---|---|
| EC→EC | 95.00 | 95.00 | 96.67 | 98.33 | 96.67 | 96.67 |
| EC→MA | 86.67 | 86.67 | 83.33 | 95.00 | 88.33 | 90.00 |
| MA→EC | 90.00 | 88.33 | 90.00 | 91.67 | 93.33 | 91.67 |
| MA→MA | 93.33 | 93.33 | 88.33 | 90.00 | 91.67 | 91.67 |
| EC→ME | 98.33 | 91.67 | 90.00 | 98.33 | 95.00 | 95.00 |
| ME→EC | 91.67 | 91.67 | 91.67 | 93.33 | 93.33 | 93.33 |
| ME→ME | 88.33 | 90.00 | 88.33 | 91.67 | 91.67 | 93.33 |
| EC→MU | 90.00 | 88.33 | 88.33 | 95.00 | 93.33 | 91.67 |
| MU→EC | 90.00 | 88.33 | 86.67 | 95.00 | 91.67 | 86.67 |
| MU→MU | 90.00 | 88.33 | 83.33 | 90.00 | 90.00 | 88.33 |

$P$: phasor; $H$: holonomy; $M$: magnitude negative-RMS; $G$: complex cosine. There are ten unique directed comparisons, not a complete 4 × 4 experiment. The shared EC→EC control is shown once.

**Supplementary Table S11:** Archived component controls: RestCog monthly EC, alpha, 6 × 50 s.

| Score | CMC@1 (%) | AUC |
|---|---|---|
| Complex cosine | 78.33 | 0.9290 |
| Complex negative-RMS | 76.67 | 0.9218 |
| Imaginary cosine | 70.00 | 0.9402 |
| Imaginary Pearson | 70.00 | 0.9394 |
| Real cosine | 60.00 | 0.8697 |
| Unit-phasor imaginary cosine | 66.67 | 0.9266 |
| Magnitude Pearson | 70.00 | 0.8827 |
| Magnitude cosine | 70.00 | 0.8831 |
| Magnitude negative-RMS | 66.67 | 0.8841 |

These values are summary-level evidence pending primary CSV import. These are separate component controls; no metric is selected retrospectively for the main comparison.

The original project archive also retains more extensive supplementary variants (for example, all cross-family fusion combinations, the optional Dortmund theta branch, and complete L-FAME duration metrics). These are archived data rather than additional manuscript figures.
